# β-adrenergic signaling modulates cancer cell mechanotype through a RhoA-ROCK-myosin II axis

**DOI:** 10.1101/777755

**Authors:** Tae-Hyung Kim, Esteban Vazquez-Hidalgo, Alexander Abdou, Xing Haw Marvin Tan, Alexei Christodoulides, Carly M. Farris, Pei-Yu Chiou, Erica K. Sloan, Parag Katira, Amy C. Rowat

## Abstract

The ability of cells to deform and generate forces are key mechanical properties that are implicated in metastasis. While various soluble and mechanical cues are known to regulate cancer cell mechanical phenotype or mechanotype, our knowledge of how cells translate external signals into changes in mechanotype is still emerging. We previously discovered that activation of β-adrenergic signaling, which results from soluble stress hormone cues, causes cancer cells to be stiffer or less deformable; this stiffer mechanotype was associated with increased cell motility and invasion. Here, we characterize how β-adrenergic activation is translated into changes in cellular mechanotype by identifying molecular mediators that regulate key components of mechanotype including cellular deformability, traction forces, and non-muscle myosin II (NMII) activity. Using a micropillar assay and computational modelling, we determine that βAR activation increases cellular force generation by increasing the number of actin-myosin binding events; this mechanism is distinct from how cells increase force production in response to matrix stiffness, suggesting that cells regulate their mechanotype using a complementary mechanism in response to stress hormone cues. To identify the molecules that modulate cellular mechanotype with βAR activation, we use a high throughput filtration platform to screen the effects of pharmacologic and genetic perturbations on βAR regulation of whole cell deformability. Our results indicate that βAR activation decreases cancer cell deformability and increases invasion by signaling through RhoA, ROCK, and NMII. Our findings establish βAR-RhoA-ROCK-NMII as a primary signaling axis that mediates cancer cell mechanotype, which provides a foundation for future interventions to stop metastasis.

## Introduction

Cellular mechanotype is implicated in multiple steps of the metastatic cascade. The ability of cancer cells to deform and generate physical forces is required during intravasation as cancer cells escape from the primary tumor (1). Indeed, cancer cells that exhibit increased traction forces tend to be more invasive (2). To metastasize, tumor cells must survive fluid shear stresses during circulation as well as mechanical stresses during their transit through narrow gaps of the vasculature (3). Cancer cell mechanotype also determines the propensity of circulating tumor cells to lodge in pulmonary beds (4, 5), which is required for seeding metastatic tumors. A variety of environmental factors are known to modulate cancer cell mechanotype, including soluble and mechanical cues. Cancer cells increase their stiffness or traction forces in response to growth factors including TGF-β (6), as well as decreased oxygen levels or hypoxia (7). Mechanical cues, such as increased matrix stiffness in tumors, cause cancer cells to increase their stiffness and force production (8–10). While various soluble and mechanical cues are known to regulate cancer cell mechanotype (11, 12), our knowledge of the shared molecular mediators that regulate cellular mechanotype and functional behaviors to drive metastasis is still emerging.

The catecholaminergic stress hormones, epinephrine and norepinephrine, are critical regulators in physiology and disease. Acute psychological stress rapidly increases local or circulating plasma concentrations of catecholamines, which results in increased activation of β-adrenoceptors (βAR); these G-protein coupled receptors are expressed in many types of cancers (13–15). We previously discovered that activation of βAR regulates the deformability of a range of cancer cell types from breast to prostate, making cells stiffer or less deformable; this stiffer mechanotype induced by βAR activation is associated with increased cell motility and invasion (16). The increased invasion of cancer cells induced by βAR activation is consistent with preclinical findings that activation of βAR signaling in response to agonists or physiological stress drives metastasis of breast cancer in mouse models (17, 18). Moreover, blocking βAR signaling with the clinically used β-blocker propranolol results in reduced metastasis and improved survival in cancer patients in prospective clinical trials (19, 20). Retrospective studies also suggest the protective effects of β-blockade as indicated by increased survival of patients who coincidentally were taking β-blockers at the time of diagnosis (21–23). If we could fully understand the molecular and biophysical mechanisms of βAR regulation of cancer cell mechanotype, this would deepen our fundamental biophysical knowledge of how cells sense and respond to environmental cues and potentially identify novel targets for intervention strategies to treat cancer.

We previously established that βAR signaling in breast cancer cells increased cell stiffness due to activation of βAR at the cell surface by soluble agonists (16). We identified that β_2_AR is the predominant subtype in highly metastatic MDA-MB-231^HM^ cells, and is required for cell stiffness changes following βAR activation (16, 24). The resultant changes in cellular deformability require both filamentous (F−) actin and non-muscle myosin II (NMII) activity (16, 25). While we established βAR regulation of cellular deformability across cancer cell types, the biophysical mechanisms and intermediate molecular mediators that translate soluble stress hormone cues into mechanical changes at the cellular level are still unclear. Here we test the hypothesis that βAR regulates the mechanotype of cancer cells through a βAR-RhoA-ROCK-NMII axis. To investigate how soluble βAR agonists regulate cellular mechanotype, we measure key components of mechanotype including cellular deformability, which is the ability of cells to deform through micron-scale pores in response to applied pressure; cellular traction forces, or the magnitude of physical forces that cells exert on their substrate; and levels of NMII activity, which are associated with cellular force production. To gain mechanistic insights into how βAR activation increases cellular traction forces, we use a computational model to determine the effects of βAR activation on the mechanisms of actomyosin-mediated force generation, including the number of NMII molecules that actively interact with actin stress fibers to generate forces. To identify molecules that mediate the βAR-induced changes in cellular mechanotype, we use a high throughput filtration platform to screen the effects of pharmacologic and genetic perturbations on whole cell deformability. Finally, we assess the role of the identified molecular mediators in cancer cell motility by measuring the *in vitro* invasion of cancer cells. Our findings establish βAR-RhoA-ROCK-NMII as a primary signaling axis that mediates cancer cell mechanotype and invasion.

## Materials and Methods

### Cell lines and reagents

The triple-negative breast adenocarcinoma cell line, MDA-MB-231, and highly metastatic variant of the cell line, MDA-MB-231^HM^, were cultured as previously described (16, 18). The non-transformed mammary epithelial cell line (MCF10A) was cultured in DMEM/F12 media supplemented with 5% horse serum, 1% pen/strep, 20 ng/ml EGF, 0.5 μg/ml hydrocortisone, 100 ng/ml cholera toxin, and 10 μg/ml insulin. The βAR agonist, isoproterenol, and antagonist, propranolol, were from MilliporeSigma (Burlington, MA, USA). To inhibit the activity of non-muscle myosin II we used (−)-blebbistatin (Selleckchem). Rho-associated protein kinases (ROCKs) were inhibited using Y-27632 dihydrochloride (Y27632, Selleckchem) and glycyl-H 1152 dihydrochloride (g-H-1152, Tocris); myosin light-chain kinase (MLCK) was inhibited using ML-7 hydrochloride (ML-7, Selleckchem); and p21-activated kinase 1 (PAK1) was inhibited using IPA-3 (Selleckchem). Cells were treated with drugs at 10 μM for 24 h prior to measurements unless stated otherwise.

### Transfections

siRNA transfections were performed using Lipofectamine 3000 (ThermoFisher) according to manufacturer’s instructions. Briefly, 50 nM of siRNAs were diluted in reduced-serum medium (Opti-MEM, Gibco) and mixed with Lipofectamine 3000 diluted in Opti-MEM followed by incubation for 5 min at room temperature. The mixture of siRNA and transfection reagent was added to the cell culture plate dropwise and cells were incubated for 72 h prior to measurement. We used the following siRNA sequences for RhoA and scrambled control (11, 26): siRhoA-#1: 5’-AUGGAAAGCAGGUAGAGUU-3’, siRhoA-#2: 5’-GAAAGACAUGCUUGCUCAU-3’, siControl: 5’-CAGUCAGGAGGAUCCAAAGTG-3’.

### Parallel microfiltration

To measure whole cell deformability, we used parallel microfiltration (PMF) (27, 28). Cells were trypsinized with 0.25% trypsin-EDTA and cells in suspension were counted using an automated cell counter (TC20, Bio-Rad) and resuspended in medium to a density of 5×10^5^ cells/ml. We also used the automated cell counter (TC20) to measure cell size distributions. To allow for cells to equilibrate after lifting into suspension, suspensions were maintained for 30 min prior to filtration. To drive cells through the 10 μm pores of the polycarbonate membrane (Millipore), we applied air pressure (2.0 kPa) for 20 s. To quantify the magnitude of cell filtration, we determined the volume of media that remained in the top well after filtration by measuring absorbance at λ_560 nm_ using a plate reader (SpectraMax M2, Molecular Devices) (29). Cells with reduced deformability have a higher probability of occluding pores and consequently exhibit a higher retention of fluid in the top well; we define the final volume of media retained in the top well compared to the initial volume loaded, Vol_final_/Vol_initial_, as % retention.

### Western blotting

Levels of proteins and protein phosphorylation were measured by western blotting. We loaded 30 μg of total protein into 4–12% Bolt gels (Invitrogen) with MES buffer (Invitrogen). Protein samples were transferred onto nitrocellulose membrane (GE Healthcare) with NuPAGE transfer buffer (Invitrogen). To minimize non-specific protein adsorption, we incubated membranes with blocking buffer (5% skim milk in TBS-T) at room temperature for 1 h. To quantify protein levels, we then incubated with the following antibodies: mouse anti-GAPDH (#MA5-15738, 1:5,000; ThermoFisher), rabbit anti-phospho-MLC2 (#3671 and #3674, 1:1,000; Cell Signaling), mouse anti-MLC2 (#4401, 1:1,000; Sigma), and mouse anti-RhoA (#MA1-134, 1:1,000; ThermoFisher). We measured the band density of the scanned film using ImageJ software (NIH, v1.50a).

### Micropillar assay

Micropillars were fabricated as previously described using PDMS and soft lithography (30). To quantify pillar dimensions, we imaged the cross-section of the micropillar mold to determine the height of 6.5 ± 0.5 μm, and imaged the top view to determine the pillar diameter of 1.75 ± 0.5 μm. To facilitate darkfield imaging with a 20× objective (NA 0.5), gold micro-disks were bonded on the top of each pillar. Prior to cell seeding, we imaged 5 regions of the pillar array. Cells were seeded and adhered overnight prior to treatment with drugs for 24 h. To delineate cells for traction force analysis, we stained cells with calcein AM (ThermoFisher) for 5 min at 37°C. The same 5 regions of the micropillar devices were then imaged using fluorescence microscopy (Zeiss Axiovert A1) equipped with a 20× objective (NA 0.5) to identify pillars occupied by cells. Darkfield microscopy was used to determine the positions of the gold-tipped pillars before and after cell seeding; displacements of the pillars that were caused by cells were determined using custom software (MATLAB). The traction force, *F*, exerted by a cell on a single pillar was determined by:

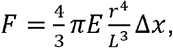

where *E* is the elastic modulus of the pillar (2.0 MPa), *r* is the radius of the pillar, *L* is the height of the pillar, and Δ*x* is the horizontal displacement of the pillar between t_0_ and t_measured_ (**Fig 1C**) (30).

**Figure 1.**
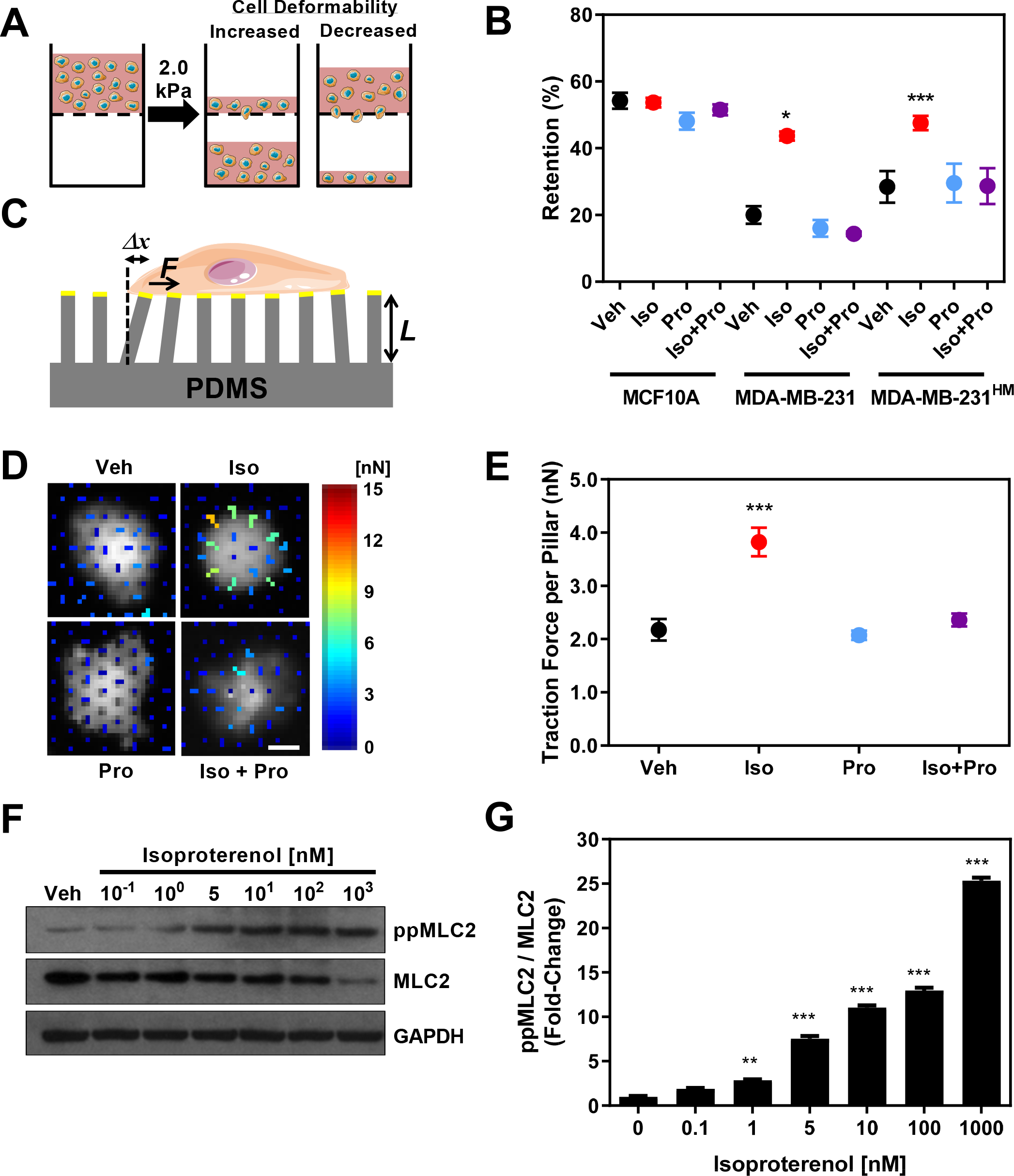
βAR activation results in decreased cancer cell deformability and increased cellular force generation. **A.** Schematic illustration showing the PMF assay. Suspensions of cells were loaded into the top well separated with porous membranes from the bottom well. **B.** Filtration measurements by PMF reveal cellular deformability of non-transformed epithelial cells (MCF10A) versus triple-negative breast cancer cells (MDA-MB-231) and the highly metastatic variant of MDA-MB-231 cells (MDA-MB-231^HM^) after treatment for 24 h with: vehicle, Veh; βAR agonist isoproterenol, Iso (100 nM); βAR antagonist propranolol, Pro (10 μM); or 100 nM Iso and 10 μM Pro. **C.** Schematic illustration showing the micropillar assay. Gold disks (yellow) were implanted on top of the pillars to facilitate imaging lateral displacements due to cellular traction forces. **D.** Representative images of MDA-MB-231^HM^ cells on micropillars with superimposed vector force map. Color scale indicates the force per pillar. Scale, 4 μm. **E.** Traction forces on individual pillars. Shown here is the median; error bars represent standard error. **F.** Western blotting against di-phosphorylated MLC2 (Thr18/Ser19), non-phosphorylated MLC2, and GAPDH. **G.** Normalized ratio of ppMLC2 to MLC2 levels after MDA-MB-231^HM^ cells were treated with increasing concentrations of isoproterenol for 2 h. Unless otherwise stated, all error bars represent mean ± s.e.m (N = 3). *P<0.05; **P<0.01; ***P<0.001 [one-way ANOVA with Tukey’s test (B, G) and Mann–Whitney test (E)]. Images in A and C are adapted from Servier Medical Art by Servier and are published under published under a Creative Commons BY license (https://creativecommons.org/licenses/by-nc/3.0/).

### Invasion assay

To measure the invasion of cells through a 3D matrix, we used a modified scratch wound assay (16, 31). A 96-well plate (ImageLock, Essen BioScience) was pre-coated with 100 μg/ml Matrigel (Corning). We then plated 3×10^4^ cells transfected with siRhoA or siControl (at post-transfection 24 h) into each well and incubated for 48 h. We generated 700-800 μm wide wounds in near 100% confluent cell monolayers using a 96-pin mechanical device (WoundMaker™, Essen BioScience). Cells were then washed with DMEM medium and 8 mg/ml Matrigel was added to cover the entire well. After a 30 m incubation at 37°C to solidify the Matrigel, 100 μl of culture medium containing isoproterenol and/or propranolol was added. We acquired images every 2 h and determined the relative wound density using IncuCyte™ software (Essen BioScience). To determine any differences in proliferation rates, which can also impact wound healing rates, we measured the proliferation of cells on Matrigel (**Supp. Fig 1B,C**).

### Computational modeling

We simulated the effect of myosin activation on cellular traction forces by modeling the transition states of myosin, the force transferred to the focal adhesion site via actin, and the bound/unbound state of integrin. Myosin has 4 transition states and a regulatory light chain dephosphorylated state 15 representing non-active myosin (**Fig 2A**). The forward transition states are *k*_*12*_, *k*_*23*_, *k*_*34*_, and *k*_*41*_. ATP hydrolysis represented by *k*_*23*_, is reversible and has a reverse rate *k*_*32*_. Dephosphorylation/phosphorylation is represented by the transition variable from state 2 to state 15, with rates *k*_*215*_ and *k*_*152*_ respectively. The ratio of transition rates between the dephosphorylated and phosphorylated myosin states are derived from the experimental ratios of ppMLC2:MLC2 obtained from western blots (**Fig 1G**),

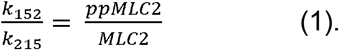

**Figure 2.**
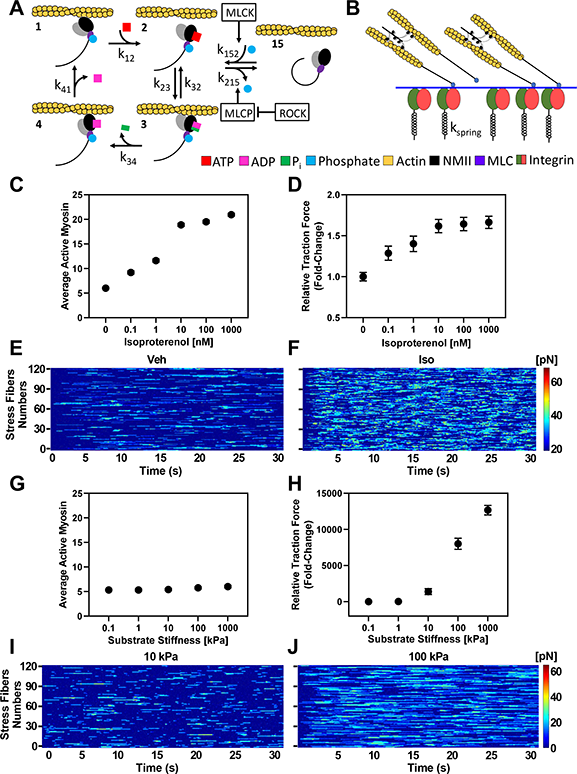
βAR activation increases the number of NMII motors interacting with F-actin to yield increased traction force generation. **A.** Model simulates the state transitions and force generation in actin stress fibers that are transferred to the substrate via integrins. **B.** Schematic shows actomyosin complex attaching to integrin on the cytosolic side. Endogenous forces affect integrin bond lifetime, which is modeled by a spring with catch-slip dynamics. **C.** Simulations predict the number of active NMII motors per actin stress fiber (for a maximum of 32 NMII per stress fiber and 120 stress fibers per μm^2^) with increasing concentrations of isoproterenol; 0 nM isoproterenol is vehicle control. **D.** Predicted traction forces per μm^2^ normalized to vehicle control. **E, F.** Kymographs show simulated forces at focal adhesions. Each row represents an individual actin stress fiber and summation over all 120 stress fibers (rows) represents the total traction force at time (*t*). Color map gradient shows magnitude of force generated at focal adhesion. **G, H.** Using the model to simulate the effect of substrate stiffness on traction forces shows effects of increasing substrate stiffness on (**G**) the number of active myosin per stress fiber and (**H**) the lifetime of catch bonds. **I, J.** Kymographs show simulated forces in individual actin stress fibers with increasing substrate stiffness (10 kPa vs 100 kPa).

Transition rates in the bound motor states 3 and 1 are modified by strain in the motor stalks due to the forces acting on the stalk (32), the effect of which is modeled as

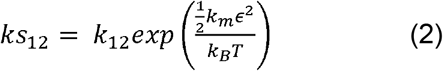

and

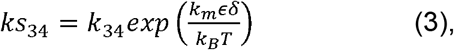

where *k*_*m*_ is the stiffness of the motor stalk, □ is the strain in the stalk, *k*_*B*_ is the Boltzmann constant, *T* is the absolute temperature, and *δ* is the characteristic bond-length of the actin-myosin bond (33). The state change of each motor is updated independently using the Gillespie algorithm to calculate the probability that a motor with change from the current state to the next by

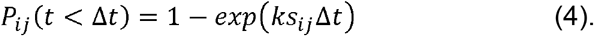

The forces generated by myosin on an actin are

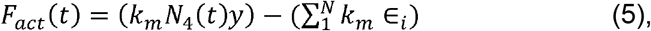

where *N*_*4*_ is the number of motors in state 4 and *y* is the motor step size. The first term represents the active force generated by the motors and the second term represents the passive force from the strain in the motor stalk of all motors (this is zero for motors that are not bound to actin filaments). The myosin-generated force on actin stress fibers is transferred to integrins at the focal adhesion/ECM interface. The displacement, *x*, of the stress-fiber pulling against the substrate is given by solution of

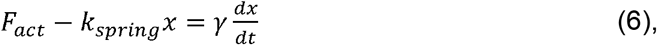

where *k*_*spring*_ is the effective spring constant of the substrate and the integrin protein, and γ is drag on the actin stress fiber (34).

The integrin catch-slip bond dissociation rate *k*_*cs*_ is modeled by

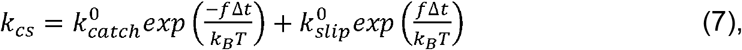

where *k*_*catch*_ represents the catch regime, *k*_*slip*_ is the slip regime, and *f* is the applied force experience by the integrin bond (35). Within a certain force range, integrin bond lifetimes increase, however, the bond will revert to a slip bond if the force exceeds the range. An unbound integrin has a probability of reattaching modeled as

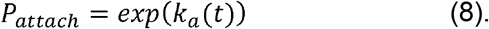

Traction forces at time (*t*) are calculated as the net force from all the actin stress fibers is obtained by

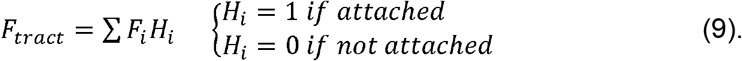

The model parameters used are obtained from existing literature and listed in supplementary table 1.

### Statistical analyses

All experiments were performed at least three independent times, unless otherwise stated. Statistical significance between control and treated groups was determined with an unpaired t-test or one-way ANOVA with Tukey’s multiple comparison post hoc analysis using GraphPad Prism 8 (GraphPad Software, La Jolla, CA).

## Results

### 1. βAR signaling modulates cancer cell mechanotype

To investigate the effect of βAR signaling on deformability of cancer cells and non-transformed epithelial cells, we used the parallel microfiltration (PMF) method that we previously developed (27). In PMF, suspensions of cells are driven to filter through a membrane with 10 μm pores on the timescale of seconds in response to applied air pressure; PMF thus measures the ability of whole cells to passively deform through narrow gaps. Less deformable cells are more likely to occlude the pores, resulting in reduced filtration and increased retention of the cell suspension in the top well (**Fig 1A**). We compared the deformability of MDA-MB-231^HM^ breast cancer cells, which is a highly metastatic variant of the established parental malignant cell line MDA-MB-231, and the non-transformed mammary epithelial cells MCF10A following treatment with the βAR agonist isoproterenol. Cells were treated with 100 nM isoproterenol, which induces maximal response in deformability (16). Both the parental MDA-MB-231 and the highly metastatic MDA-MB-231^HM^ cells showed similar increases in retention with isoproterenol treatment (**Fig 1B**). By contrast we found no significant changes in the retention of MCF10A cells following βAR activation. We did not observe any significant changes in cell size across treatment conditions (**Supp. Fig 1A**), indicating that the observed differences in retention with isoproterenol treatment were not due to changes in cell size. We also found that the increased retention caused by isoproterenol was abrogated by treatment with the βAR antagonist propranolol, confirming that isoproterenol is acting through βAR. These findings suggest that βAR activation alters the deformability of malignant cancer cells.

Since we previously showed that the increased stiffness of breast cancer cells with activation of βAR signaling was dependent on NMII activity (16, 25), and NMII contributes to actomyosin-mediated cellular force generation, we speculated that βAR may regulate cellular traction forces. NMII activity is also critical for cancer cell invasion (36) and more invasive cancer cells tend to have increased traction forces compared to less invasive cells (2). To test the hypothesis that activation of βAR signaling increases cellular force generation, we treated MDA-MB-231^HM^ cells with βAR agonist and measured cellular traction forces using a micropillar assay. Cells were plated on flexible polydimethylsiloxane (PDMS) micropillars, and the lateral displacements of pillars were tracked after activation of βAR with the agonist isoproterenol (**Fig 1C**) (30). Activation of βAR signaling in MDA-MB-231^HM^ cells with 100 nM of isoproterenol resulted in a ~2-fold increase in median traction forces from ~2 nN to ~4 nN per pillar (p < 0.0001) (**Fig 1E**); this increase in cellular traction forces was abrogated by the β-blocker propranolol. Propranolol itself had no effects on baseline cellular traction forces compared to vehicle (**Fig 1D, E**). These findings show that βAR activation increases cancer cell force generation.

To determine how βAR activation impacts the activity of NMII, we measured phosphorylation of myosin light chain 2 (MLC2), which regulates NMII motor activity, the formation of myosin filaments, and actin-myosin crosslinking (36). We found that βAR activation by isoproterenol resulted in a concentration-dependent increase in the phosphorylation of MLC2 (**Fig 1F, G**). With the same isoproterenol concentration that induced an increase in cellular traction forces, we observed a ~13-fold increase in MLC2 phosphorylation relative to total MLC2. Taken together, these results indicate that βAR activation increases NMII activity and cellular traction forces.

### 2. βAR activation increases cellular force generation by enhancing actin-NMII binding

To gain mechanistic insight into how stress hormones regulate cellular mechanotype, we generated a computational model to predict cellular traction forces. In this model, focal adhesions were modeled as multiple actin stress fibers (~ 120/μm^2^) (37–39) tugging on substrate bound integrins by the action of myosin motors (~ 32/stress fiber) (40, 41). Individual motor-filament interactions as well as integrin dynamics were simulated using a stochastic Monte Carlo approach (42). The activity of each individual motor was a function of its phosphorylation state, ATP binding and hydrolysis rates, as well as the pushing or pulling forces acting on each motor (**Fig 2A, Supp. Table 1**). We used the experimentally determined ratio of ppMLC2 to total MLC2 (**Fig 1G**) as an input to the model to capture ROCK-dependent MLC2 phosphorylation and dephosphorylation rates. Using this model, we estimated the force in individual actin stress fibers over time as they bind, tug, and unbind from surface bound integrins. We also determined the net traction force per unit area generated within focal adhesions over time. Increasing isoproterenol concentration from 0 to 1,000 nM resulted in a 4-fold increase in the number of NMII molecules that actively interact with actin filaments and generate forces (**Fig 2C**). Consequently, for isoproterenol concentrations > 10 nM there is a ~2-fold increase in the predicted net traction force per unit area (**Fig 2D**), in close quantitative agreement with the experimental traction force data (**Fig 1E**). Interestingly, the increased number of interacting NMII molecules in response to βAR signaling does not significantly increase the force generated within each individual stress fiber, but rather the total number of stress fibers generating traction forces greater than 20 pN within a focal adhesion as shown in the kymographs (**Fig 2E, F, Supp. Fig 2A, B**). To examine how this mechanism of cellular force generation compares with the cellular response to matrix stiffness, which is established to increase cellular traction forces (43, 44), we used the same model to predict traction forces with increasing substrate stiffness (increasing the value of effective *k*_*spring*_ in the model). We observed that with increasing substrate stiffness, the model predicts increased traction force at the focal adhesion results from a stronger force generated within a single stress fiber while the number of interacting motors remains the same (**Fig 2G-J**); this is due to an increase in the average lifetime of integrin catch bonds on stiffer substrates, allowing the forces in the stress fibers to reach higher values before the bond breaks (**Supp. Fig 2D-F**). Here we have neglected downstream effects of focal adhesion signaling in modeling the response to cells to both βAR activation and matrix stiffness. However, the results obtained from our minimal model sufficiently capture our experimental observations of βAR activation on cellular force generation (**Fig 1E**), indicating that we are capturing the predominant mechanisms of traction force generation. Moreover, our observations are consistent with previous models of cellular traction force generation with increasing substrate stiffness (45). Taken together, these data from both computational modeling (**Fig 2**) and experiments (**Fig 1E**) substantiate that βAR activation increases cellular force generation by increasing the number of active motors per stress fiber and suggest that cells may tune force production through distinct yet complementary mechanisms for soluble versus mechanical cues.

### 3. βAR signaling alters cell deformability through a RhoA-ROCK-NMII axis

To begin to dissect the molecular mechanisms underlying how βAR increases NMII activity to regulate cellular mechanotype, we tested the role of specific kinases that may be involved in βAR regulation of NMII activity using pharmacologic inhibitors. We activated βAR signaling while simultaneously inhibiting the activity of three kinases that are well-characterized regulators of NMII activity: Rho-associated protein kinase (ROCK), myosin light-chain kinase (MLCK), and p21-activated kinase (PAK) using the pharmacologic inhibitors Y-27632, ML-7, and IPA-3. We used PMF to rapidly assay effects of these perturbations on βAR regulation of cellular deformability (27). Activating βAR by isoproterenol treatment resulted in a significant increase in retention (as in **Fig 1B**). However, when NMII activity was inhibited with blebbistatin prior to isoproterenol treatment, we observed no significant changes in retention (**Fig 3A**). Similarly, when ROCK was inhibited by Y-27632, isoproterenol treatment did not cause any significant increase in retention, suggesting that ROCK is also required for the βAR-induced decrease in cellular deformability. By contrast, when MLCK and PAK were inhibited by ML-7 and IPA-3, we still observed a statistically significant increase in retention following isoproterenol treatment (**Fig 3A**). Taken together, these findings indicate that ROCK is a major contributor to the βAR-induced changes in cellular deformability. To further validate that ROCK is an essential kinase in βAR-regulation of cell deformability, we treated cells with increasing concentrations of isoproterenol in the presence of each myosin kinase inhibitor (**Fig 3B-D**). The βAR-induced increase in retention was fully blocked only when ROCK activity was inhibited by Y27632 (**Fig 3B**). We observed similar effects with the ROCK inhibitor, g-H-1152, which has higher specificity for ROCK than Y-27632 (46) (**Supp. Fig 1E)**. These results show that ROCK activity is required for βAR regulation of cellular deformability.

**Figure 3.**
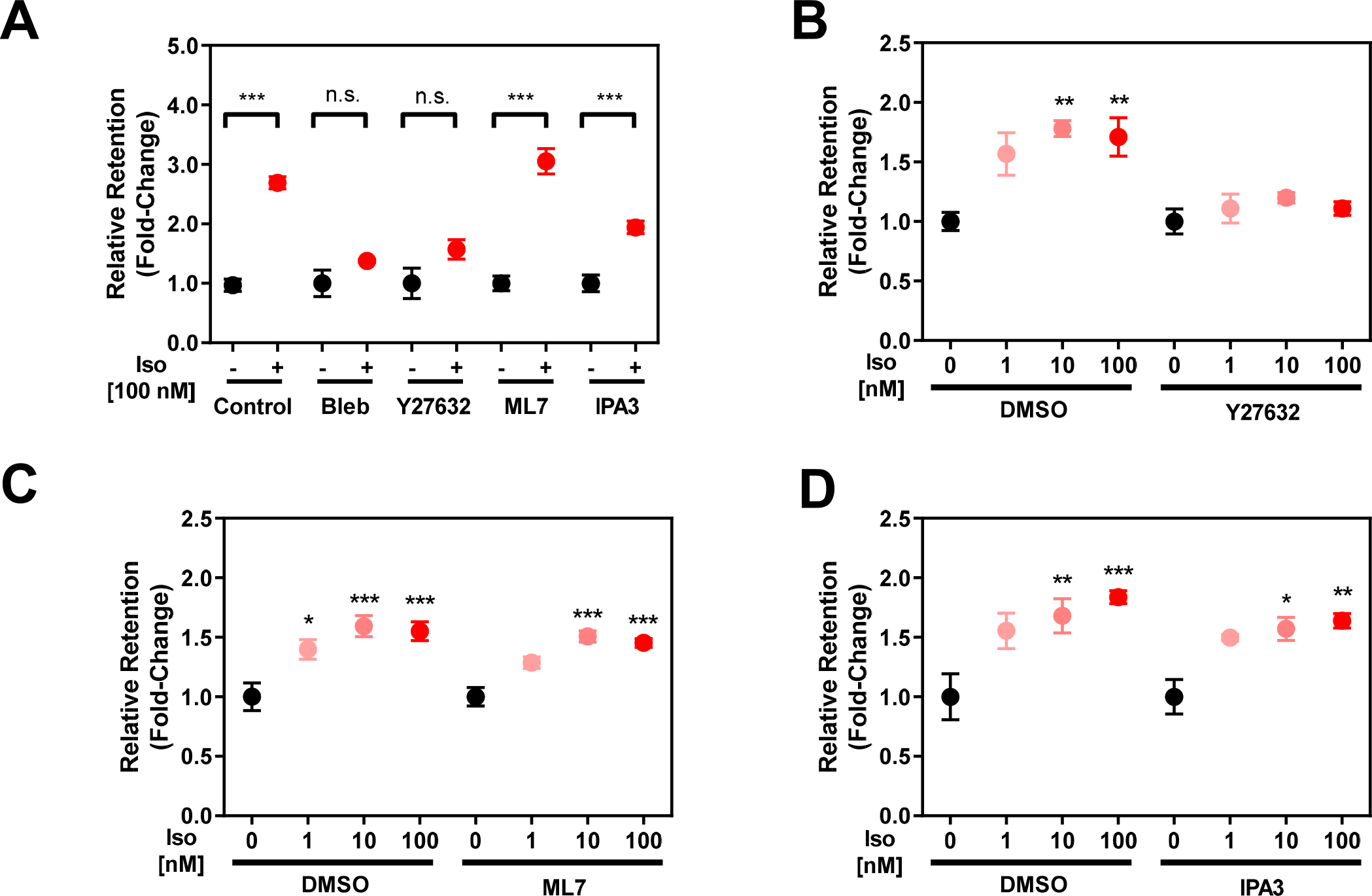
βAR-induced changes in cellular deformability require ROCK activity. **A.** Cell filtration measurements by PMF with simultaneous βAR activation by isoproterenol (Iso) and suppression of myosin activity by pharmacological inhibitors, Bleb, blebbistatin (10 μM); Y27632 (10 μM); ML7 (10 μM); and IPA3 (10 μM). Cells were co-treated with inhibitors and isoproterenol for 24 h prior to filtration measurements. **B-D.** Filtration measurements with increasing concentration of isoproterenol with or without myosin light chain kinase inhibitors. n.s.: not significant, *P<0.05; **P<0.01; ***P<0.001 [one-way ANOVA with Tukey’s test].

We next investigated the role of RhoA, a canonical upstream regulator of ROCK and NMII activity. To address the role of RhoA in regulating βAR-induced changes in cellular deformability, we knocked down RhoA using two different siRNAs (siRhoA-1 and siRhoA-2), and assayed cell filtration following isoproterenol treatment. Knockdown of RhoA by siRNA reduced protein levels by ~70% (**Fig 4A, B**). Cells with RhoA knockdown did not show any observable increase in retention following βAR activation (**Fig 4C**); these findings were consistent with the effects of ROCK inhibition (**Fig 3A, B, Supp. Fig 1E**) and support that βAR activation alters cellular deformability through a RhoA-ROCK-NMII axis.

**Figure 4.**
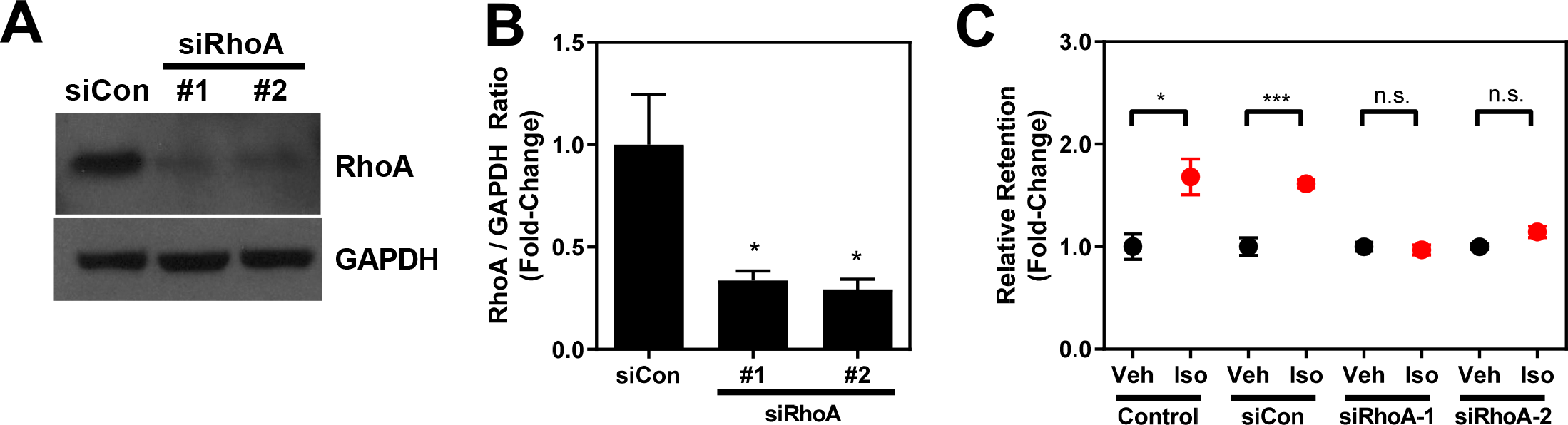
RhoA contributes to βAR regulation of cellular deformability. **A.** Confirmation of siRNA knockdown of RhoA by western blotting. GAPDH is loading control. **B.** Ratio of RhoA to GAPDH normalized to control siRNA (siCon). **C.** Cell filtration with PMF after 100 nM of isoproterenol treatment for 24 h in control and RhoA knockdown cells. n.s.: not significant, *P<0.05; ***P<0.001 [one-way ANOVA with Tukey’s test].

### 4. βAR signaling regulates cell invasion through a RhoA-ROCK-NMII axis

Since cellular mechanotype is strongly associated with the motility of cancer cells (16, 47–49), we next investigated the role of a RhoA-ROCK-NMII axis in regulating the βAR-induced changes in the invasion of breast cancer cells. To simulate invasion through the tissue environment, we used a 3D scratch wound assay (16, 31). We found that isoproterenol treatment increased invasion compared to vehicle treated cells (**Fig 5A, C**); this finding was consistent with our previous studies (16). However, with inhibition of NMII activity (blebbistatin) or ROCK (Y-27632), βAR activation had no observable effects on cell invasion (**Fig 5A, C, E**). We next tested the effects of RhoA on βAR modulation of cancer cell invasion. We found that RhoA knockdown also suppressed the βAR-mediated increase in cell invasion (**Fig 5B, D, F**). Since cell proliferation can also impact wound closure rates in this 3D invasion assay, we measured changes in cell confluence over the experimental timescale of 48 h but found no significant changes across inhibitor treatments and RhoA knockdowns (**Supp. Fig 1B, C**). These findings demonstrate that the increased invasion due to βAR activation requires ROCK, RhoA, and NMII activity.

**Figure 5.**
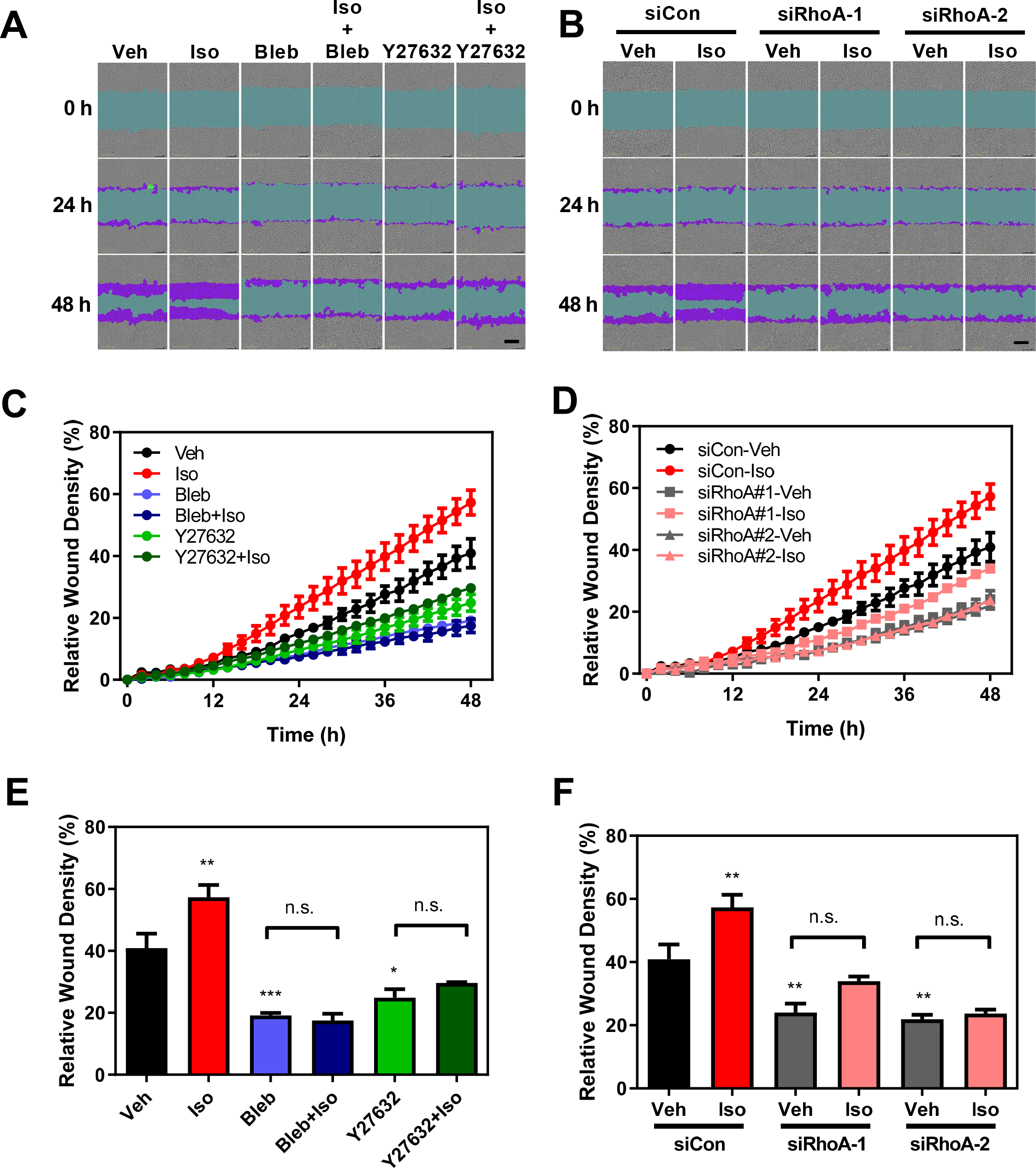
NMII, ROCK, and RhoA activity are required for the increased cell invasion due to βAR activation. **A, B.** Representative images from a 3D scratch wound invasion assay. Drugs were added at time 0: isoproterenol (Iso, 100 nM), NMII inhibitor blebbistatin (Bleb, 10 μM), and ROCK inhibitor Y27632 (10 μM). Transfections were performed 72 h prior to time 0 h of the invasion assay. Scale: 300 μm. **C, D.** Relative wound density as a function of time. **E, F**. Relative wound density at 48 h. Unless otherwise indicated, all comparisons were made to vehicle control (Veh). n.s.: not significant, *P<0.05; **P<0.01; ***P<0.001 [one-way ANOVA with Tukey’s test].

Taken together, our results show that a βAR-RhoA-ROCK-NMII axis plays a central role in how cancer cells translate soluble stress hormone cues into changes in cellular mechanotype, including their deformability and contractility, as well as invasion (**Fig 6**). Mechanistically our findings show that the altered mechanotype that occurs with βAR activation results from increased NMII-actin interactions.

**Figure 6.**
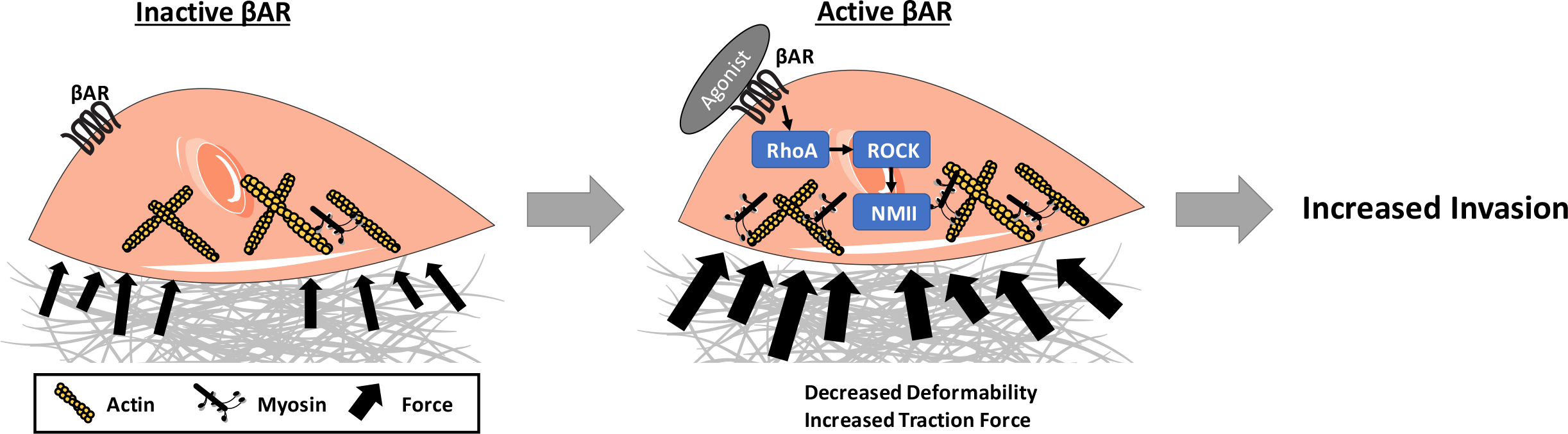
**Schematic illustration of the proposed mechanism for how a βAR-RhoA-ROCK-NMII axis regulates cancer cell mechanotype** and functional behaviors that contribute to metastasis. Actin stress finbers are shown in yellow and myosins attached to stress fibers are shown in black. Black arrows indicate traction forces generated by the cell. Images are adapted from Servier Medical Art by Servier (http://www.servier.com/Powerpoint-image-bank) and are published under published under a Creative Commons BY license (https://creativecommons.org/licenses/by-nc/3.0/).

## Discussion

Here we show that a soluble βAR agonist elicits mechanotype changes in cancer cells through a βAR-RhoA-ROCK-NMII axis. Specifically, we found that cancer cells exhibit increased NMII activity and increased number of NMII-actin interactions, as well as enhanced traction force generation in response to βAR activation, and that RhoA, ROCK, and NMII activity contribute to the decreased deformability and increased invasion of cancer cells with βAR activation (**Fig 6**).

By developing a mechanistic model that predicts the mechanotype changes induced by βAR activation, we gained a deeper understanding of how cells translate soluble signals into mechanical forces. Our findings reveal that βAR activation increases cellular force generation by increasing the number of NMII motors engaged with actin filaments and in turn increasing the number of actin stress fibers generating forces at a given time; this is driven by ROCK dependent changes in MLC phosphorylation states. We showed how this βAR-regulated mechanism of increased cellular force production contrasts the mechanism of how cells enhance their force production in response to increased matrix stiffness: modeling stiffer substrates we observed an increase in cellular traction forces due to increased forces generated by individual actin stress fibers, which results from an increased lifetime of integrin-substrate catch bonds. The ability of cells to increase force production through two separate yet complementary mechanisms could enable them to independently tune force production to achieve enhanced sensitivity and/or dynamic range in response to combinations of external cues. Our results that show βAR activation increases the number of actin-bound NMII motors by ~4-fold, which also explains the decreased whole cell deformability that we observed using the PMF assay. Existing models show that cell stiffness increases linearly with NMII motor-generated stress in biopolymer networks and this motor-generated stress is in turn proportional to the number of active force generating motors in the system (50, 51). The computational models we generated here can be further extended to predict the effects of soluble cues on more complex cellular behaviors such as motility and bi-directional cell-matrix interactions. Future work will determine how simultaneous matrix stiffness and stress hormone cues are processed and integrated to regulate cellular force generation and mechanical homeostasis. More broadly, the framework that we have established could be applied to advance knowledge of how combinations of diverse types of external cues impact cellular mechanotype. The approach we present here shows how such a combined experimental and computational framework is critical to address the fundamental question of how cells process diverse soluble and mechanical cues to generate an integrated cellular response.

Here we establish βAR-RhoA-ROCK-NMII as a mechanoregulating pathway that may be used by cancer cells to regulate their deformability, traction force generation, and invasion. Both changes in mechanotype and invasion can impact tumorigenesis and metastasis (1, 52). While RhoA and ROCK are well established regulators of cellular motility and cancer cell invasion (53, 54), knowledge of the upstream signals that activate this pathway is still emerging. The RhoA-ROCK axis has been implicated in how cancer cells sense and respond to the increased matrix stiffness that results from the increased tumor stiffness due to fibrosis and cancer associated fibroblast contractility (55); this results in integrin signaling through a RhoA-ROCK axis (56). More recently, hypoxia has been shown to alter cancer cell force generation through RhoA-ROCK signaling (7). Other essential factors in the tumor microenvironment such as epidermal growth factors have been shown to modulate cancer cell mechanotype including local cell elasticity, actin cytoskeleton architecture, and cell migration (12). It will be interesting in future work to define the interplay among multiple signals that activate RhoA-ROCK (57) to regulate cancer cell behaviors in the complex tumor microenvironment.

The molecular mediators upstream of RhoA that determine βAR-mediated regulation of mechanotype also remain to be determined. βAR is known to regulate cellular functions through the canonical G protein-mediated cAMP-PKA signaling pathway (58–60). Our previous findings support that βAR-regulation of cellular mechanotype involves cAMP-PKA because treatment with the adenyl cyclase activator forskolin also resulted in similar decreased cellular deformability (16). In addition to the cAMP-PKA pathway, βAR could regulate cellular mechanotype through a βArrestin-Src pathway (61, 62). The role of specific ROCK isoforms in βAR-regulation of mechanotype also remains to be determined, including whether the two isoforms of the ROCK, ROCK1 and ROCK2, play redundant roles. Previous reports identify that ROCK isoforms differentially modulate cancer cell motility in MDA-MB-231 cells (56), so we anticipate that specific isoforms may also differentially modulate cellular mechanotype. While we show here RhoA-ROCK is a major axis for activating NMII through βAR, additional pathways that regulate NMII activity, such as MLCK and PAK, may be also be involved in βAR-regulation of cancer cell mechanotype through compensatory effects and/or crosstalk with the RhoA-ROCK signaling pathway (63, 64). Understanding the interplay among diverse signaling pathways that regulate cancer cell invasion and metastasis will also be crucial; for example, βAR activation affects downstream mediators including Src and MMPs, which are established to promote metastasis (61, 62, 65, 66). Combinatorial studies that utilize co-treatments of inhibitors and/or knockdowns will further refine our knowledge of the specific pathways that contribute to βAR regulation of cellular mechanotype.

Given the role of epithelial cells in regulating morphogenesis (67, 68), wound healing (69), and cancer progression (70), a deeper knowledge of how stress hormones impact cellular mechanotype should advance our understanding of physiological processes. Future studies across a broader range of malignant and non-transformed epithelial cell types will be valuable to define how broadly the regulation of cellular mechanotype through a βAR-RhoA-ROCK-NMII axis holds across different types of cells. Our findings from this study suggest that RhoA-ROCK is the primary mechanism of βAR-regulation of breast cancer cell mechanotype. βAR signaling is established to increase the contractility of cardiac myocytes (71) and to decrease the traction stresses of human airway smooth muscle cells (72). Recent findings show that activating βAR in adipose cells results in increased NMII activity and cell contractility, albeit through a Ca^2+^-MLCK pathway (73). Taken together, our observations contribute to the growing literature that shows βAR activation regulates force generation across a range of cell types through slightly different mechanisms; such findings suggest that βAR regulation of cellular mechanotype is a convergent phenomenon that is essential to cellular homeostasis. For cancer cells, βAR-induced mechanotype changes could reflect their altered invasive behavior, which could provide them with a selective advantage to metastasize. Future *in vivo* studies will help to elucidate the role of cellular mechanotype in βAR regulation of metastasis. Ultimately a deeper understanding of the molecular signaling pathways that regulate cancer cell behaviors will advance our knowledge of cancer biophysics and benefit the rational design of more effective drugs to suppress metastasis.

## Author Contributions

T-H.K., P-Y.C, E.K.S. and A.C.R. designed the experiments. T-H.K., A.A., X.H.T., E.K.S., and A.C. performed experiments and analyzed the data. E.V-H., C.M.F., P.K. designed and conducted the computational modeling. E.V-H., P.K., T-H.K., A.C.R. contributed to the iterative design of experiments and simulations. T-H.K., E.K.S., P.K., and A.C.R. wrote the manuscript. All authors edited and approved the final manuscript.

## Acknowledgments

We thank the Flow Cytometry Core at the Broad Stem Cell Research Center at UCLA for the IncuCyte^®^. We also thank UCLA Statistical Consulting Group for their statistical assistance.

This work was supported by grants from the National Science Foundation (BMMB-1906165 to A.C.R. and BMMB-1905390, BMMB-1763132 to P.K.), the Army Research Office (W911NF-17-1-0413 to P.K.), the Jonsson Comprehensive Cancer Foundation, and the University of California Cancer Research Coordinating Committee (CRR-18-526901 to A.C.R.).

The authors declare there are no conflicts of interest.

---

**Supplementary Figure 1**

**A.** Cell sizes were measured after drug treatment with isoproterenol (Iso, 100 nM), propranolol (Pro, 10 μM), and co-treatment with Iso and Pro. **B, C.** Cell proliferation rates were measured after drug treatment with Iso, blebbistatin (Bleb, 10 μM), the ROCK inhibitor Y27632 (10 μM), as well as co-treatment with inhibitors and isoproterenol, including after RhoA knockdown. The dashed line indicates the time point (48 h) at which cell invasion was measured in **Fig 5E, F**. To assess statistical significance at 48 h, we performed one-way ANOVA with Tukey’s multiple comparison post hoc analysis between control and treated groups for (B) and an unpaired t-test between Veh and Iso for each siRNA for (C). **D.** Cell size measurements after treatment with g-H-1152 and Y27632 with or without Iso. **E.** Filtration measurements with increasing concentration of isoproterenol with or without g-H-1152 (1 μM). All filtration data is reported as mean±s.e.m. Boxplots show the lower and upper quartiles with median (line), with the whiskers representing the 10–90th percentiles. Individual dots represent outliers. n.s.: not significant, *P<0.05; **P<0.01; ***P<0.001 [one-way ANOVA with Tukey’s test].

**Supplementary Figure 2**

**A.** The maximum force per filament, **B.** number of individual stress fibers attached to focal adhesions, and **C.** bond lifetime as a function of isoproterenol concentration. **D.** The maximum force per filament, **E.** number of individual stress fibers attached to focal adhesions, and **F.** bond lifetime with increasing substrate stiffness.

## References

1. Wirtz, D., K. Konstantopoulos, and P.C. Searson. 2011. The physics of cancer: the role of physical interactions and mechanical forces in metastasis. Nat. Rev. Cancer. 11: 512–522.

2. Kraning-Rush, C.M., J.P. Califano, and C.A. Reinhart-King. 2012. Cellular traction stresses increase with increasing metastatic potential. PLoS One. 7: e32572.

3. Barnes, J.M., J.T. Nauseef, and M.D. Henry. 2012. Resistance to fluid shear stress is a conserved biophysical property of malignant cells. PLoS One. 7: e50973.

4. Shaw Bagnall, J., S. Byun, S. Begum, D.T. Miyamoto, V.C. Hecht, S. Maheswaran, S.L. Stott, M. Toner, R.O. Hynes, and S.R. Manalis. 2015. Deformability of Tumor Cells versus Blood Cells. Sci. Rep. 5: 18542.

5. Harouaka, R.A., M. Nisic, and S.-Y. Zheng. 2013. Circulating tumor cell enrichment based on physical properties. J Lab Autom. 18: 455–468.

6. Gladilin, E., S. Ohse, M. Boerries, H. Busch, C. Xu, M. Schneider, M. Meister, and R. Eils. 2019. TGFβ-induced cytoskeletal remodeling mediates elevation of cell stiffness and invasiveness in NSCLC. Sci. Rep. 9: 7667.

7. Gilkes, D.M., L. Xiang, S.J. Lee, P. Chaturvedi, M.E. Hubbi, D. Wirtz, and G.L. Semenza. 2014. Hypoxia-inducible factors mediate coordinated RhoA-ROCK1 expression and signaling in breast cancer cells. Proc. Natl. Acad. Sci. USA. 111: E384–93.

8. Trichet, L., J. Le Digabel, R.J. Hawkins, S.R.K. Vedula, M. Gupta, C. Ribrault, P. Hersen, R. Voituriez, and B. Ladoux. 2012. Evidence of a large-scale mechanosensing mechanism for cellular adaptation to substrate stiffness. Proc. Natl. Acad. Sci. USA. 109: 6933–6938.

9. Yeung, T., P.C. Georges, L.A. Flanagan, B. Marg, M. Ortiz, M. Funaki, N. Zahir, W. Ming, V. Weaver, and P.A. Janmey. 2005. Effects of substrate stiffness on cell morphology, cytoskeletal structure, and adhesion. Cell Motil. Cytoskeleton. 60: 24–34.

10. Solon, J., I. Levental, K. Sengupta, P.C. Georges, and P.A. Janmey. 2007. Fibroblast adaptation and stiffness matching to soft elastic substrates. Biophys. J. 93: 4453–4461.

11. Ingallina, E., G. Sorrentino, R. Bertolio, K. Lisek, A. Zannini, L. Azzolin, L.U. Severino, D. Scaini, M. Mano, F. Mantovani, A. Rosato, S. Bicciato, S. Piccolo, and G. Del Sal. 2018. Mechanical cues control mutant p53 stability through a mevalonate-RhoA axis. Nat. Cell Biol. 20: 28–35.

12. Azadi, S., M. Tafazzoli-Shadpour, M. Soleimani, and M.E. Warkiani. 2019. Modulating cancer cell mechanics and actin cytoskeleton structure by chemical and mechanical stimulations. J. Biomed. Mater. Res. 107: 1569–1581.

13. Wan, C., C. Gong, H. Zhang, L. Hua, X. Li, X. Chen, Y. Chen, X. Ding, S. He, W. Cao, Y. Wang, S. Fan, Y. Xiao, G. Zhou, and A. Shen. 2016. β2-adrenergic receptor signaling promotes pancreatic ductal adenocarcinoma (PDAC) progression through facilitating PCBP2-dependent c-myc expression. Cancer Lett. 373: 67–76.

14. Qin, J., F. Jin, N. Li, H. Guan, L. Lan, H. Ni, and Y. Wang. 2015. Adrenergic receptor β2 activation by stress promotes breast cancer progression through macrophages M2 polarization in tumor microenvironment. BMB Rep. 48: 295–300.

15. Braadland, P.R., H. Ramberg, H.H. Grytli, and K.A. Taskén. 2014. β-Adrenergic Receptor Signaling in Prostate Cancer. Front. Oncol. 4: 375.

16. Kim, T.-H., N.K. Gill, K.D. Nyberg, A.V. Nguyen, S.V. Hohlbauch, N.A. Geisse, C.J. Nowell, E.K. Sloan, and A.C. Rowat. 2016. Cancer cells become less deformable and more invasive with activation of β-adrenergic signaling. J. Cell Sci. 129: 4563–4575.

17. Sloan, E.K., S.J. Priceman, B.F. Cox, S. Yu, M.A. Pimentel, V. Tangkanangnukul, J.M.G. Arevalo, K. Morizono, B.D.W. Karanikolas, L. Wu, A.K. Sood, and S.W. Cole. 2010. The sympathetic nervous system induces a metastatic switch in primary breast cancer. Cancer Res. 70: 7042–7052.

18. Le, C.P., C.J. Nowell, C. Kim-Fuchs, E. Botteri, J.G. Hiller, H. Ismail, M.A. Pimentel, M.G. Chai, T. Karnezis, N. Rotmensz, G. Renne, S. Gandini, C.W. Pouton, D. Ferrari, A. Möller, S.A. Stacker, and E.K. Sloan. 2016. Chronic stress in mice remodels lymph vasculature to promote tumour cell dissemination. Nat. Commun. 7: 10634.

19. Shaashua, L., M. Shabat-Simon, R. Haldar, P. Matzner, O. Zmora, M. Shabtai, E. Sharon, T. Allweis, I. Barshack, L. Hayman, J. Arevalo, J. Ma, M. Horowitz, S. Cole, and S. Ben-Eliyahu. 2017. Perioperative COX-2 and β-Adrenergic Blockade Improves Metastatic Biomarkers in Breast Cancer Patients in a Phase-II Randomized Trial. Clin. Cancer Res. 23: 4651–4661.

20. De Giorgi, V., M. Grazzini, S. Benemei, N. Marchionni, E. Botteri, E. Pennacchioli, P. Geppetti, and S. Gandini. 2018. Propranolol for Off-label Treatment of Patients With Melanoma: Results From a Cohort Study. JAMA Oncol. 4: e172908.

21. Barron, T.I., R.M. Connolly, L. Sharp, K. Bennett, and K. Visvanathan. 2011. Beta blockers and breast cancer mortality: a population- based study. J. Clin. Oncol. 29: 2635–2644.

22. Melhem-Bertrandt, A., M. Chavez-Macgregor, X. Lei, E.N. Brown, R.T. Lee, F. Meric-Bernstam, A.K. Sood, S.D. Conzen, G.N. Hortobagyi, and A.-M. Gonzalez-Angulo. 2011. Beta-blocker use is associated with improved relapse-free survival in patients with triple-negative breast cancer. J. Clin. Oncol. 29: 2645–2652.

23. Powe, D.G., M.J. Voss, K.S. Zänker, H.O. Habashy, A.R. Green, I.O. Ellis, and F. Entschladen. 2010. Beta-blocker drug therapy reduces secondary cancer formation in breast cancer and improves cancer specific survival. Oncotarget. 1: 628–638.

24. Pon, C.K., J.R. Lane, E.K. Sloan, and M.L. Halls. 2016. The β2-adrenoceptor activates a positive cAMP-calcium feedforward loop to drive breast cancer cell invasion. FASEB J. 30: 1144–1154.

25. Kim, T.-H., C. Ly, A. Christodoulides, C.J. Nowell, P.W. Gunning, E.K. Sloan, and A.C. Rowat. 2019. Stress hormone signaling through β-adrenergic receptors regulates macrophage mechanotype and function. FASEB J. 33: 3997–4006.

26. Pillé, J.Y., C. Denoyelle, J. Varet, J.R. Bertrand, J. Soria, P. Opolon, H. Lu, L.L. Pritchard, J.P. Vannier, C. Malvy, C. Soria, and H. Li. 2005. Anti-RhoA and anti-RhoC siRNAs inhibit the proliferation and invasiveness of MDA-MB-231 breast cancer cells in vitro and in vivo. Mol. Ther. 11: 267–274.

27. Qi, D., N. Kaur Gill, C. Santiskulvong, J. Sifuentes, O. Dorigo, J. Rao, B. Taylor-Harding, W. Ruprecht Wiedemeyer, and A.C. Rowat. 2015. Screening cell mechanotype by parallel microfiltration. Sci. Rep. 5: 17595.

28. Gill, N.K., N.K. Gill, D. Qi, T.-H. Kim, C. Chan, A. Nguyen, K.D. Nyberg, and A.C. Rowat. 2017. A protocol for screening cells based on deformability using parallel microfiltration. Protoc exch..

29. Gill, N.K., C. Ly, K.D. Nyberg, L. Lee, D. Qi, B. Tofig, M. Reis-Sobreiro, O. Dorigo, J. Rao, R. Wiedemeyer, B. Karlan, K. Lawrenson, M.R. Freeman, R. Damoiseaux, and A.C. Rowat. 2019. A scalable filtration method for high throughput screening based on cell deformability. Lab Chip. 19: 343–357.

30. Xiao, F., X. Wen, X.H.M. Tan, and P.-Y. Chiou. 2018. Plasmonic micropillars for precision cell force measurement across a large field-of-view. Appl. Phys. Lett. 112: 033701.

31. Pan, Y., G. Robertson, L. Pedersen, E. Lim, A. Hernandez-Herrera, A.C. Rowat, S.L. Patil, C.K. Chan, Y. Wen, X. Zhang, U. Basu-Roy, A. Mansukhani, A. Chu, P. Sipahimalani, R. Bowlby, D. Brooks, N. Thiessen, C. Coarfa, Y. Ma, R.A. Moore, J.E. Schein, A.J. Mungall, J. Liu, C.V. Pecot, A.K. Sood, S.J.M. Jones, M.A. Marra, and P.H. Gunaratne. 2016. miR-509-3p is clinically significant and strongly attenuates cellular migration and multi-cellular spheroids in ovarian cancer. Oncotarget. 7: 25930–25948.

32. Marcucci, L., and C. Reggiani. 2016. Mechanosensing in Myosin Filament Solves a 60 Years Old Conflict in Skeletal Muscle Modeling between High Power Output and Slow Rise in Tension. Front. Physiol. 7: 427.

33. Stam, S., J. Alberts, M.L. Gardel, and E. Munro. 2015. Isoforms confer characteristic force generation and mechanosensation by myosin II filaments. Biophys. J. 108: 1997–2006.

34. Colombelli, J., A. Besser, H. Kress, E.G. Reynaud, P. Girard, E. Caussinus, U. Haselmann, J.V. Small, U.S. Schwarz, and E.H.K. Stelzer. 2009. Mechanosensing in actin stress fibers revealed by a close correlation between force and protein localization. J. Cell Sci. 122: 1665–1679.

35. Rakshit, S., and S. Sivasankar. 2014. Biomechanics of cell adhesion: how force regulates the lifetime of adhesive bonds at the single molecule level. Phys. Chem. Chem. Phys. 16: 2211–2223.

36. Vicente-Manzanares, M., X. Ma, R.S. Adelstein, and A.R. Horwitz. 2009. Non-muscle myosin II takes centre stage in cell adhesion and migration. Nat. Rev. Mol. Cell Biol. 10: 778–790.

37. Young, L.E., and H.N. Higgs. 2018. Focal Adhesions Undergo Longitudinal Splitting into Fixed-Width Units. Curr. Biol. 28: 2033–2045.e5.

38. Hu, S., Y.-H. Tee, A. Kabla, R. Zaidel-Bar, A. Bershadsky, and P. Hersen. 2015. Structured illumination microscopy reveals focal adhesions are composed of linear subunits. Cytoskeleton (Hoboken). 72: 235–245.

39. Cramer, L.P., M. Siebert, and T.J. Mitchison. 1997. Identification of novel graded polarity actin filament bundles in locomoting heart fibroblasts: implications for the generation of motile force. J. Cell Biol. 136: 1287–1305.

40. Mak, M., M.H. Zaman, R.D. Kamm, and T. Kim. 2016. Interplay of active processes modulates tension and drives phase transition in self-renewing, motor-driven cytoskeletal networks. Nat. Commun. 7: 10323.

41. Miller, C.J., D. Harris, R. Weaver, G.B. Ermentrout, and L.A. Davidson. 2018. Emergent mechanics of actomyosin drive punctuated contractions and shape network morphology in the cell cortex. PLoS Comput. Biol. 14: e1006344.

42. Farris, C.M. 2017. The Role of Myosin Head State and Binding Site Availability in Calcium-Dependent Regulation of a Muscle Mimetic System. Masters Abstracts International. 57: 91.

43. Califano, J.P., and C.A. Reinhart-King. 2010. Substrate stiffness and cell area predict cellular traction stresses in single cells and cells in contact. Cell Mol. Bioeng. 3: 68–75.

44. Han, S.J., K.S. Bielawski, L.H. Ting, M.L. Rodriguez, and N.J. Sniadecki. 2012. Decoupling substrate stiffness, spread area, and micropost density: a close spatial relationship between traction forces and focal adhesions. Biophys. J. 103: 640–648.

45. Marzban, B., X. Yi, and H. Yuan. 2018. A minimal mechanics model for mechanosensing of substrate rigidity gradient in durotaxis. Biomech Model Mechanobiol. 17: 915–922.

46. Tamura, M., H. Nakao, H. Yoshizaki, M. Shiratsuchi, H. Shigyo, H. Yamada, T. Ozawa, J. Totsuka, and H. Hidaka. 2005. Development of specific Rho-kinase inhibitors and their clinical application. Biochim. Biophys. Acta. 1754: 245–252.

47. Swaminathan, V., K. Mythreye, E.T. O’Brien, A. Berchuck, G.C. Blobe, and R. Superfine. 2011. Mechanical stiffness grades metastatic potential in patient tumor cells and in cancer cell lines. Cancer Res. 71: 5075–5080.

48. Xu, W., R. Mezencev, B. Kim, L. Wang, J. McDonald, and T. Sulchek. 2012. Cell stiffness is a biomarker of the metastatic potential of ovarian cancer cells. PLoS One. 7: e46609.

49. Nyberg, K.D., S.L. Bruce, A.V. Nguyen, C.K. Chan, N.K. Gill, T.-H. Kim, E.K. Sloan, and A.C. Rowat. 2018. Predicting cancer cell invasion by single-cell physical phenotyping. Integr Biol (Camb). 10: 218–231.

50. Broedersz, C.P., and F.C. MacKintosh. 2011. Molecular motors stiffen non-affine semiflexible polymer networks. Soft Matter. 7: 3186.

51. Chen, P., and V.B. Shenoy. 2011. Strain stiffening induced by molecular motors in active crosslinked biopolymer networks. Soft Matter. 7: 355–358.

52. Swift, J., I.L. Ivanovska, A. Buxboim, T. Harada, P.C.D.P. Dingal, J. Pinter, J.D. Pajerowski, K.R. Spinler, J.-W. Shin, M. Tewari, F. Rehfeldt, D.W. Speicher, and D.E. Discher. 2013. Nuclear lamin-A scales with tissue stiffness and enhances matrix-directed differentiation. Science. 341: 1240104.

53. Yoshioka, K., S. Nakamori, and K. Itoh. 1999. Overexpression of small GTP-binding protein RhoA promotes invasion of tumor cells. Cancer Res. 59: 2004–2010.

54. Yamazaki, D., S. Kurisu, and T. Takenawa. 2009. Involvement of Rac and Rho signaling in cancer cell motility in 3D substrates. Oncogene. 28: 1570–1583.

55. Vennin, C., V.T. Chin, S.C. Warren, M.C. Lucas, D. Herrmann, A. Magenau, P. Melenec, S.N. Walters, G. Del Monte-Nieto, J.R.W. Conway, M. Nobis, A.H. Allam, R.A. McCloy, N. Currey, M. Pinese, A. Boulghourjian, A. Zaratzian, A.A.S. Adam, C. Heu, A.M. Nagrial, A. Chou, A. Steinmann, A. Drury, D. Froio, M. Giry-Laterriere, N.L.E. Harris, T. Phan, R. Jain, W. Weninger, E.J. McGhee, R. Whan, A.L. Johns, J.S. Samra, L. Chantrill, A.J. Gill, M. Kohonen-Corish, R.P. Harvey, A.V. Biankin, Australian Pancreatic Cancer Genome Initiative (APGI), T.R.J. Evans, K.I. Anderson, S.T. Grey, C.J. Ormandy, D. Gallego-Ortega, Y. Wang, M.S. Samuel, O.J. Sansom, A. Burgess, T.R. Cox, J.P. Morton, M. Pajic, and P. Timpson. 2017. Transient tissue priming via ROCK inhibition uncouples pancreatic cancer progression, sensitivity to chemotherapy, and metastasis. Sci. Transl. Med. 9: eaai8504.

56. Peng, Y., Z. Chen, Y. Chen, S. Li, Y. Jiang, H. Yang, C. Wu, F. You, C. Zheng, J. Zhu, Y. Tan, X. Qin, and Y. Liu. 2019. ROCK isoforms differentially modulate cancer cell motility by mechanosensing the substrate stiffness. Acta Biomater. 88: 86–101.

57. Daubriac, J., S. Han, J. Grahovac, E. Smith, A. Hosein, M. Buchanan, M. Basik, and Y. Boucher. 2018. The crosstalk between breast carcinoma-associated fibroblasts and cancer cells promotes RhoA-dependent invasion via IGF-1 and PAI-1. Oncotarget. 9: 10375–10387.

58. Carr, R., J. Schilling, J. Song, R.L. Carter, Y. Du, S.M. Yoo, C.J. Traynham, W.J. Koch, J.Y. Cheung, D.G. Tilley, and J.L. Benovic. 2016. β-arrestin-biased signaling through the β2-adrenergic receptor promotes cardiomyocyte contraction. Proc. Natl. Acad. Sci. USA. 113: E4107–16.

59. Barnes, W.G., E. Reiter, J.D. Violin, X.-R. Ren, G. Milligan, and R.J. Lefkowitz. 2005. beta-Arrestin 1 and Galphaq/11 coordinately activate RhoA and stress fiber formation following receptor stimulation. J. Biol. Chem. 280: 8041–8050.

60. Limbird, L.E., D.M. Gill, and R.J. Lefkowitz. 1980. Agonist-promoted coupling of the beta-adrenergic receptor with the guanine nucleotide regulatory protein of the adenylate cyclase system. Proc. Natl. Acad. Sci. USA. 77: 775–779.

61. Luttrell, L.M., S.S. Ferguson, Y. Daaka, W.E. Miller, S. Maudsley, G.J. Della Rocca, F. Lin, H. Kawakatsu, K. Owada, D.K. Luttrell, M.G. Caron, and R.J. Lefkowitz. 1999. Beta-arrestin-dependent formation of beta2 adrenergic receptor-Src protein kinase complexes. Science. 283: 655–661.

62. Armaiz-Pena, G.N., J.K. Allen, A. Cruz, R.L. Stone, A.M. Nick, Y.G. Lin, L.Y. Han, L.S. Mangala, G.J. Villares, P. Vivas-Mejia, C. Rodriguez-Aguayo, A.S. Nagaraja, K.M. Gharpure, Z. Wu, R.D. English, K.V. Soman, M.M.K. Shahzad, M. Zigler, M.T. Deavers, A. Zien, T.G. Soldatos, D.B. Jackson, J.E. Wiktorowicz, M. Torres-Lugo, T. Young, K. De Geest, G.E. Gallick, M. Bar-Eli, G. Lopez-Berestein, S.W. Cole, G.E. Lopez, S.K. Lutgendorf, and A.K. Sood. 2013. Src activation by β-adrenoreceptors is a key switch for tumour metastasis. Nat. Commun. 4: 1403.

63. Kassianidou, E., J.H. Hughes, and S. Kumar. 2017. Activation of ROCK and MLCK tunes regional stress fiber formation and mechanics via preferential myosin light chain phosphorylation. Mol. Biol. Cell. 28: 3832–3843.

64. Zhang, W., B.P. Bhetwal, and S.J. Gunst. 2018. Rho kinase collaborates with p21-activated kinase to regulate actin polymerization and contraction in airway smooth muscle. J. Physiol. (Lond.). 596: 3617–3635.

65. Barbieri, A., S. Bimonte, G. Palma, A. Luciano, D. Rea, A. Giudice, G. Scognamiglio, E. La Mantia, R. Franco, S. Perdonà, O. De Cobelli, M. Ferro, S. Zappavigna, P. Stiuso, M. Caraglia, and C. Arra. 2015. The stress hormone norepinephrine increases migration of prostate cancer cells in vitro and in vivo. Int. J. Oncol. 47: 527–534.

66. He, Q., G. Wu, and M.C. Lapointe. 2000. Isoproterenol and cAMP regulation of the human brain natriuretic peptide gene involves Src and Rac. Am. J. Physiol. Endocrinol. Metab. 278: E1115–23.

67. Nedvetsky, P.I., S.-H. Kwon, J. Debnath, and K.E. Mostov. 2012. Cyclic AMP regulates formation of mammary epithelial acini in vitro. Mol. Biol. Cell. 23: 2973–2981.

68. Wei, S.C., L. Fattet, J.H. Tsai, Y. Guo, V.H. Pai, H.E. Majeski, A.C. Chen, R.L. Sah, S.S. Taylor, A.J. Engler, and J. Yang. 2015. Matrix stiffness drives epithelial-mesenchymal transition and tumour metastasis through a TWIST1-G3BP2 mechanotransduction pathway. Nat. Cell Biol. 17: 678–688.

69. Handly, L.N., and R. Wollman. 2017. Wound-induced Ca2+ wave propagates through a simple release and diffusion mechanism. Mol. Biol. Cell. 28: 1457–1466.

70. Liu, X., J. Li, B.L. Cadilha, A. Markota, C. Voigt, Z. Huang, P.P. Lin, D.D. Wang, J. Dai, G. Kranz, A. Krandick, D. Libl, H. Zitzelsberger, I. Zagorski, H. Braselmann, M. Pan, S. Zhu, Y. Huang, S. Niedermeyer, C.A. Reichel, B. Uhl, D. Briukhovetska, J. Suárez, S. Kobold, O. Gires, and H. Wang. 2019. Epithelial-type systemic breast carcinoma cells with a restricted mesenchymal transition are a major source of metastasis. Sci. Adv. 5: eaav4275.

71. Ribeiro, A.J.S., O. Schwab, M.A. Mandegar, Y.-S. Ang, B.R. Conklin, D. Srivastava, and B.L. Pruitt. 2017. Multi-Imaging Method to Assay the Contractile Mechanical Output of Micropatterned Human iPSC-Derived Cardiac Myocytes. Circ. Res. 120: 1572–1583.

72. Wang, N., I.M. Tolić-Nørrelykke, J. Chen, S.M. Mijailovich, J.P. Butler, J.J. Fredberg, and D. Stamenović. 2002. Cell prestress. I. Stiffness and prestress are closely associated in adherent contractile cells. Am. J. Physiol. Cell Physiol. 282: C606–16.

73. Tharp, K.M., M.S. Kang, G.A. Timblin, J. Dempersmier, G.E. Dempsey, P.-J.H. Zushin, J. Benavides, C. Choi, C.X. Li, A.K. Jha, S. Kajimura, K.E. Healy, H.S. Sul, K. Saijo, S. Kumar, and A. Stahl. 2018. Actomyosin-Mediated Tension Orchestrates Uncoupled Respiration in Adipose Tissues. Cell Metab. 27: 602–615.e4.

