## Supplemental Table 1 for "β-adrenergic signaling modulates cancer cell mechanotype through a RhoA-ROCK-myosin II axis"

| Parameter | Definition | Value | Unit | Reference |
| --- | --- | --- | --- | --- |
| $\Delta t$ | Monte Carlo simulation time step | 1 | ms |  |
| k_12_ | Transition rate from state 1 to 2 | 1.4x10^-4^ | ms^-1^ | [1] |
| k_23_ | Transition rate from state 2 to 3 | 0.007 | ms^-1^ | [1] |
| k_32_ | Reverse transition rate from state 3 to 2 | 0.011 | ms^-1^ | [1] |
| k_34_ | Transition rate from state 3 to 4 | 1.6x10^-4^ | ms^-1^ | [1] |
| k_41_ | Transition rate from state 4 to 1 | 0.028 | ms^-1^ | [1] |
| k_152_ | Phosphorylation rate | 1 | ms^-1^ | Estimated |
| k_215_ | Dephosphorylation rate | k_152_ x $\frac{MLC}{ppMLC}$ | ms^-1^ | Calculated |
| k_B_ | Boltzman constant | 1.3806x10^-2^ | J/K |  |
| k_spring_ | Effective substrate-integrin spring constant | 1-1000 | pN/nm |  |
| E | Substrate Stiffness | Related to k_spring_, E = k_spring_/a, where a is a characteristic distance of substrate material |  | [3] |
| T | Temperature | 300 | K |  |
| δ | Distance between myosin binding sites | 2.5 | nm | [2] |
| k_m_ | Stiffness of motor stalk | 4 | pN/nm | [3] |
| y | Motor step size | 5.3 | nm | [4] |
| k^0^_catch_ | Integrin catch rate | 0.055 | ms^-1^ | [5] |
| k^0^_slip_ | Integrin slip rate | 5.2x10^-7^ | ms^-1^ | [5] |
| k­_a_ | Integrin attach rate | 1x10^2^ | ms^-1^ | [5] |

**Supplementary Table 1.** Model parameters.

[1] Kovacs M, Wang F, Hu A, Zhang Y, Sellers JR. Functional divergence of human cytoplasmic myosin II. The Journal of Biological Chemistry. 2003. 278(40):38132-38140.
[2] Stam S, Alberts J, Gardel ML, Munro E. Isoforms confer characteristic force generation and mechanosensation by myosin II filaments. Biophysical Journal. 2015. 108(8):1997-2006
[3] Howard, J. Mechanics of Motor Proteins and the Cytoskeleton. 1^st^ ed. Sunderland, MA. Sinauer Associates. 2001,
[4] Kitamua K, Tokunaga M, Iwane AH, Yanaia T. A single myosin head moves along an actin filament with regular steps of 5.3 nanometres. Nature. 397:129-134. 1999
[5] Rakshit, S., and S. Sivasankar. 2014. Biomechanics of cell adhesion: how force regulates the lifetime of adhesive bonds at the single molecule level. Phys. Chem. Chem. Phys. 16: 2211–2223
