## Supplementary figures and images for "β-adrenergic signaling modulates cancer cell mechanotype through a RhoA-ROCK-myosin II axis"

### Supplemental Figures

#### Supplementary Figure 1

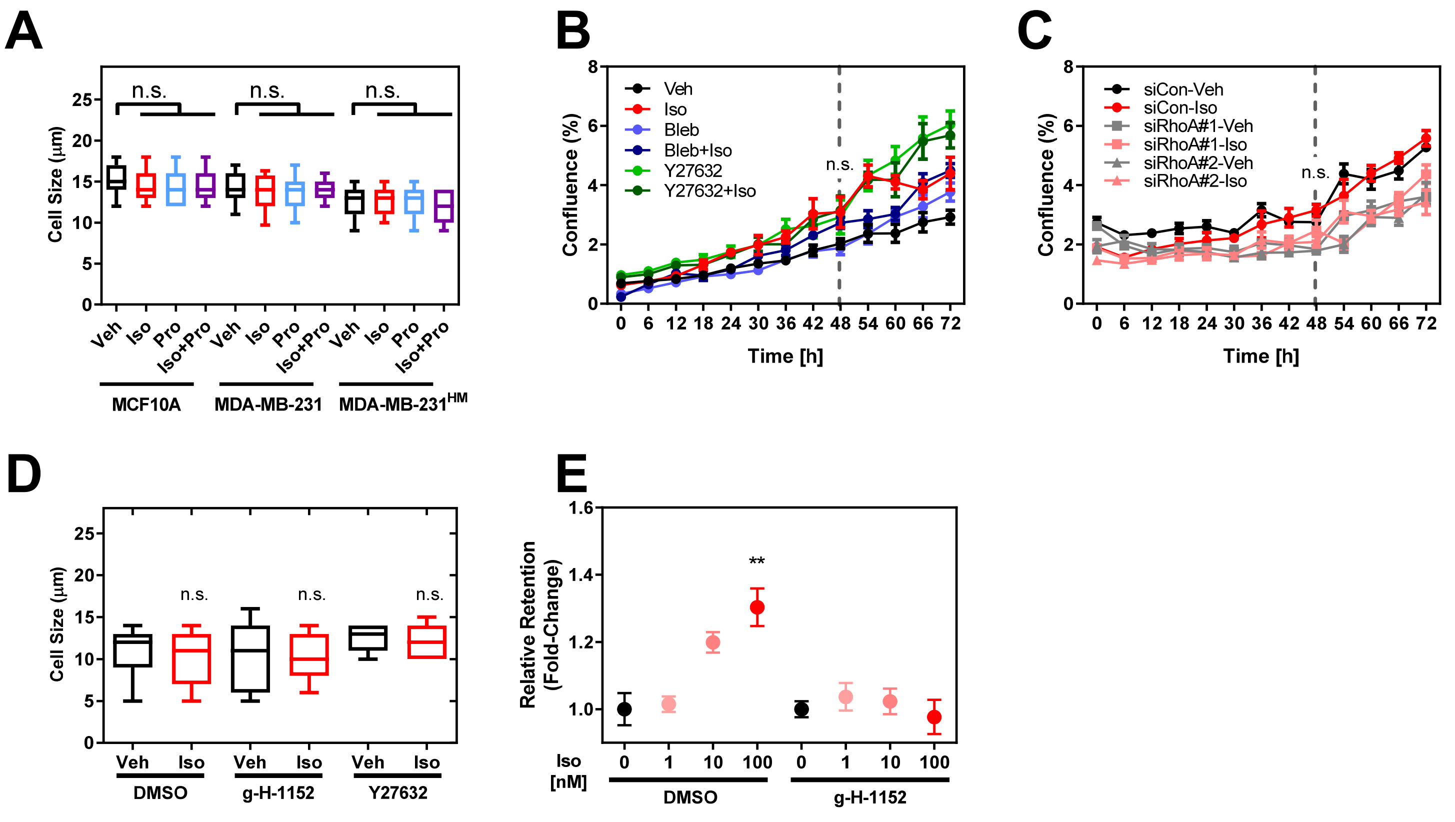

#### Supplementary Figure 2

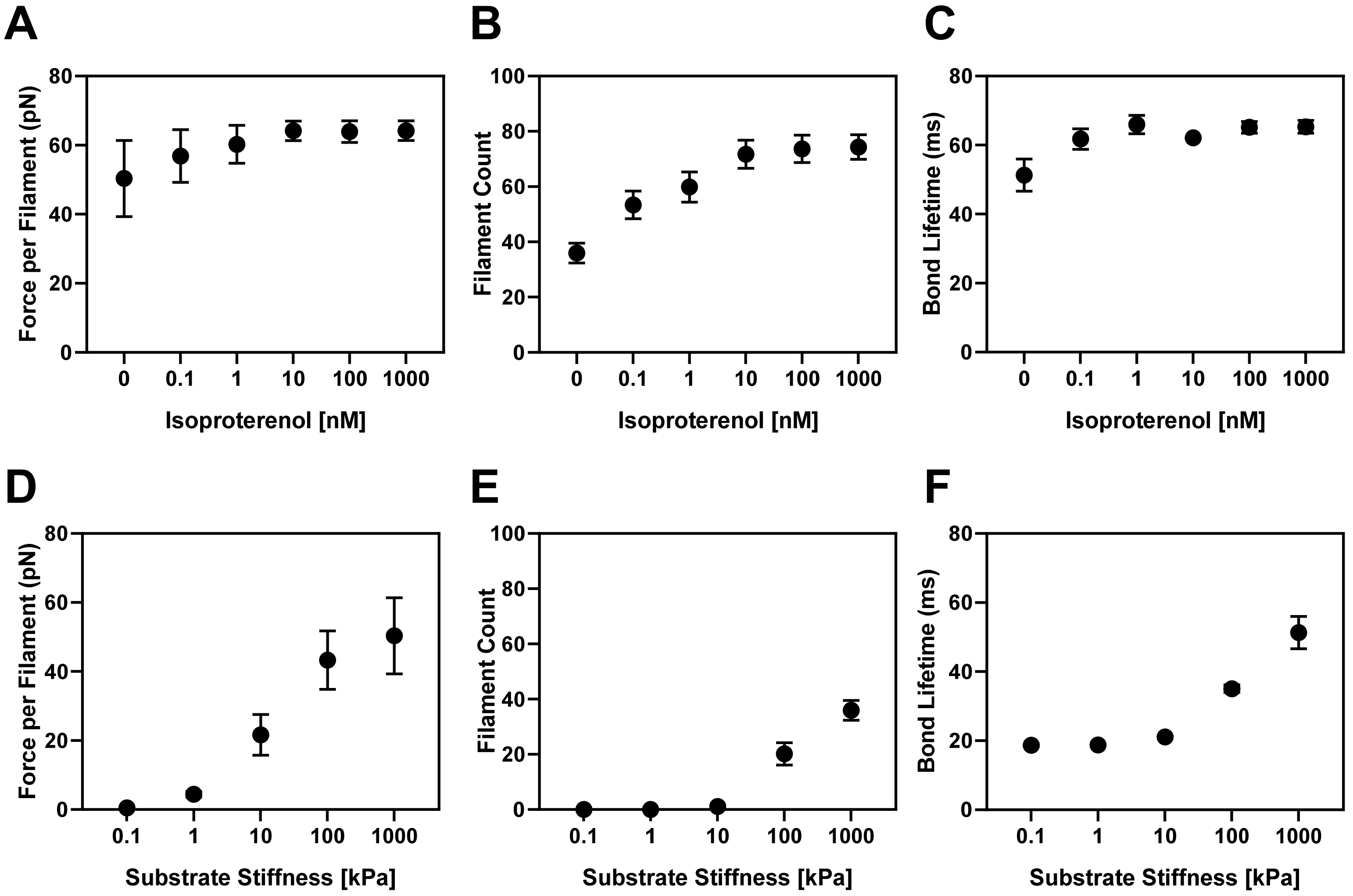
